# Ensemble projections of global ocean animal biomass with climate change

**DOI:** 10.1101/467175

**Authors:** Heike K. Lotze, Derek P. Tittensor, Andrea Bryndum-Buchholz, Tyler D. Eddy, William W. L. Cheung, Eric D. Galbraith, Manuel Barange, Nicolas Barrier, Daniele Bianchi, Julia L. Blanchard, Laurent Bopp, Matthias Büchner, Catherine Bulman, David A. Carozza, Villy Christensen, Marta Coll, John Dunne, Elizabeth A. Fulton, Simon Jennings, Miranda Jones, Steve Mackinson, Olivier Maury, Susa Niiranen, Ricardo OliverosRamos, Tilla Roy, José A. Fernandes, Jacob Schewe, Yunne-Jai Shin, Tiago A. M. Silva, Jeroen Steenbeek, Charles A. Stock, Philippe Verley, Jan Volkholz, Nicola D. Walker

## Abstract

Climate change is shifting the abundance and distribution of marine species with consequences for ecosystem functioning, seafood supply, management and conservation. Several approaches for future projection exist but these have never been compared systematically to assess their variability. We conducted standardized ensemble projections including 6 global fisheries and marine ecosystem models, forced with 2 Earth-system models and 4 emission scenarios in a fished and unfished ocean, to derive average trends and associated uncertainties. Without fishing, mean global animal biomass decreased by 5% (±4%) under low and 17% (±11%) under high emissions by 2100, primarily driven by increasing temperature and decreasing primary production. These climate-change effects were slightly weaker for larger animals and in a fished ocean. Considerable regional variation ranged from strong biomass increases in high latitudes to strong decreases in mid-low latitudes, with good model agreement on the direction of change but variable magnitude. Uncertainties due to differences among ecosystem or Earth-system models were similar, suggesting equal need for model improvement. Our ensemble projections provide the most comprehensive outlook on potential climate-driven ecological changes in the ocean to date. Realized future trends will largely depend on how fisheries and management adapt to these changes in a changing climate.

## Introduction

Climate change is shifting the abundance and distribution of marine species^1-3^ with consequences for ecosystem functioning, seafood supply, management and conservation^4-6^. Quantifying future trends in natural ecosystems due to climate change, including estimates of uncertainties is critical to inform ongoing climate-change and biodiversity assessments (IPCC, IPBES)^3^ and guide pathways towards achieving key policy objectives, such as the UN Sustainable Development Goals (SDGs). Various modeling approaches are used to assess potential impacts^6-10^, yet each model is a necessary but inherently incomplete simplification of the natural world, often built on heterogeneous assumptions and incorporating different processes^11^. One approach to overcome individual-model limitations is to force a range of models with standardized climate-change scenarios and combine them into ensemble projections to estimate mean future trends and associated inter-model spread^11^. Such Model Inter-comparison Projects (MIPs) have proven critical for enhancing credibility and understanding of future climate-change projections^12^ and impacts on Earth’s freshwater, vegetation and agriculture^13-15^, but have not yet been attempted for the global marine ecosystem.

Over the past decade, several global fisheries and marine ecosystem models (EMs) have been developed, characterized by different scopes, structures and assumptions^11^. Some of these have been used individually to project future changes in species distribution, biomass or potential fisheries catch^6-10^, but it remains unclear how consistent and comparable individual model results are, and thus how applicable for providing robust insight and advice.

We assessed projected changes in global marine animal biomass over the 21^st^ century through ensemble projections with six published EMs (Table 1, Table S1) forced with standardized outputs from two Earth-system models (ESMs) and four emission scenarios (Representative Concentration Pathways, RCPs). The ESMs spanned the range of available CMIP5 projections^12^, from low (GFDL-ESM2M) to high (IPSL-CM5A-LR) increases in sea surface temperature (SST) and associated changes in net primary production (NPP, Fig. S1), while other drivers were more similar^12^. The climate-change scenarios were run for a historical (1970-2005) and future period (2006-2100) without fishing to isolate the climate signal, and with fishing to evaluate how climate-driven responses differ in a fished ocean. Future fishing drivers were held constant at 2005 levels because temporally- and spatially-explicit future fishing scenarios are not yet available^11^. All EMs reported standardized outputs of total animal biomass and biomass of animals >10cm and >30cm. Not all EMs could run the full set of scenarios due to EM or ESM limitations, so we analyzed all available runs for each scenario, and performed sensitivity analyses on subsets, which revealed qualitatively and quantitatively similar results (Table S2).

**Table 1.** Overview on the six global marine fisheries and ecosystem models used in ensemble projections. See Table S1 for more detail.

| **Model name** | **Basic structure** | **Taxonomic scope** | **Key ecological processes** |
| --- | --- | --- | --- |
| BiOeconomic mArine Trophic Size-spectrum Model (BOATS) | Size structure | 3 size groups representing all commercial fish | Metabolic rates, biomass flow, empirical mortality, interactive fishing effort |
| Macroecological Model | Size structure | 180 body mass classes | Metabolic rates, prey size selection, predation efficiency |
| Dynamic Pelagic Benthic Model (DPBM) | Size structure, trait-based | 1 pelagic predator and 1 benthic detritivore size spectrum with 100 size classes each | Size-based predation (pelagic) and competition (benthic) driving growth and mortality, vital rates |
| Dynamic Bioclimate Envelope Model (DBEM) | Species distribution | 892 commercial fish and invertebrate species | Carrying capacity as a function of habitat suitability, vital rates, movement of adults and larval dispersal |
| EcoOcean Model | Trophodynamic | 51 trophic groups including invertebrates and vertebrates | Biomass and production flow, predation, competition, foraging capacity, movement of trophic groups by dispersal |
| Apex Predators ECOSystem Model (APECOSM) | 3D-, trait- and size-structured communities | 3 communities (epipelagic, migratory, mesopelagic) with 95 species length classes and 100 size classes | Size-based predation, competition, metabolic rates, 3D passive transport, 3D active movement, schooling |

## Results and Discussion

Our ensemble projections revealed consistent declines in global marine animal biomass from 1970-2100. Without fishing, mean declines in total biomass ranged from 4.8% (±3.5% standard deviation) under low (RCP2.6) to 17.2% (±10.7%) under high (RCP8.5) emissions by 2090-2099 relative to 1990-1999 (Fig. 1a). All four emission scenarios projected similar declines in total biomass by 2030, the target year of many SDGs, and through to mid-century, after which they began to diverge. Projected mean declines were similar for animals >10cm and >30cm (Fig. 1c, Fig. S2), albeit slightly lower and more variable for animals >30cm (Table S2). Thus, the consequences of different emission scenarios may not be distinguishable over the next 2-3 decades but should differ markedly in the long term.

**Figure 1.**
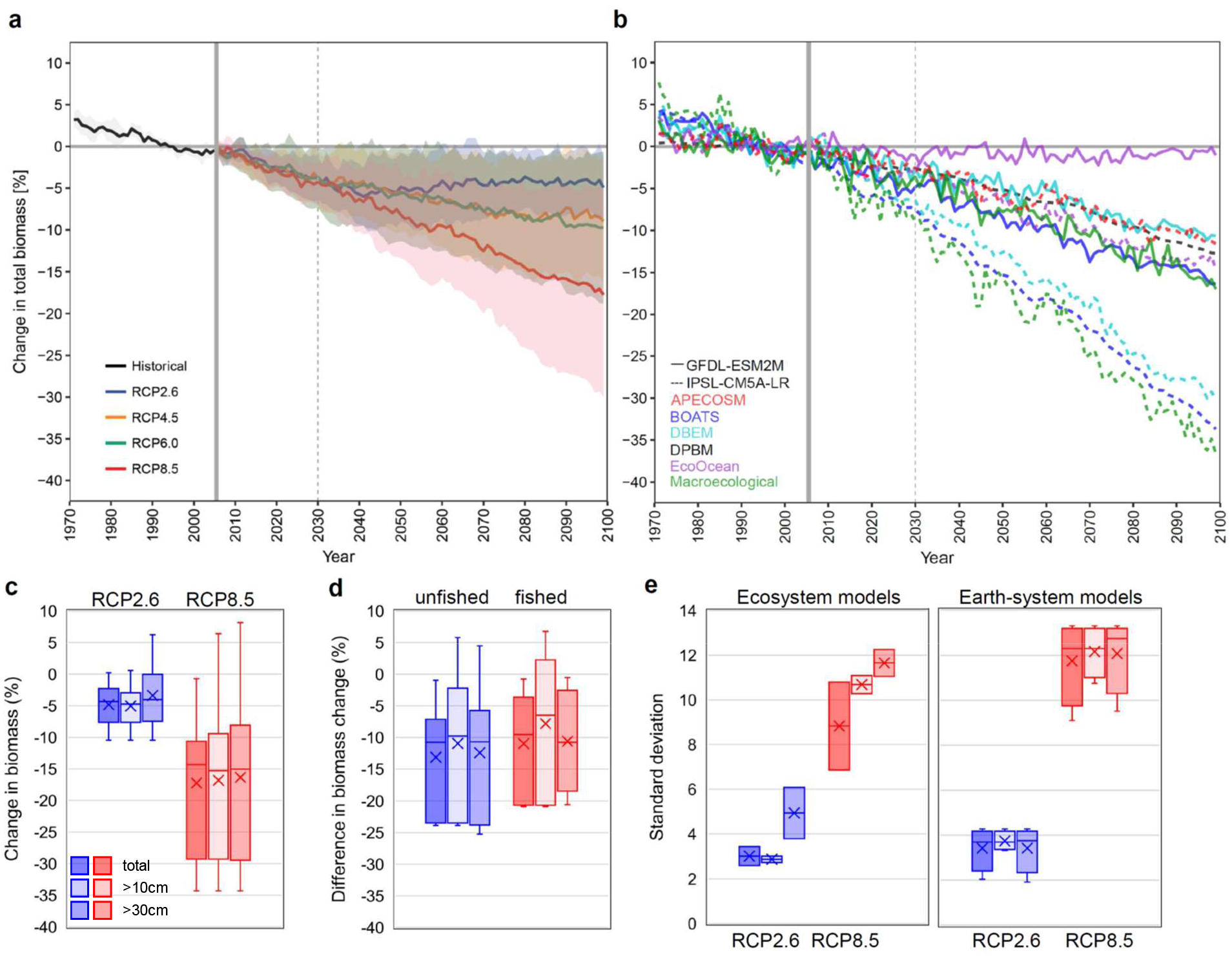
Ensemble projections of global ocean animal biomass with climate change. (**a**) Multi-model mean change in total biomass without fishing for historical and 4 future emission scenarios (RCPs) relative to 1990-99 with one standard deviation (±SD, n=10); thin vertical line indicates the target year for achieving the Sustainable Development Goals. (**b**) Individual model projections for RCP8.5 without fishing showing the spread across different ecosystem and Earth-system model combinations. (**c**) Projected biomass declines for different size groups in 2090-99 relative to 1990-99 under low (RCP2.6) and high (RCP8.5) emissions without fishing (n=10). (**d**) Projected climate-change effect (RCP8.5 vs. 2.6) in 2090-99 for different size groups in an unfished compared to a fished ocean (n=6). (**e**) Variability in projected biomass declines (expressed as % standard deviation) due to the use of different Earth-system models and different ecosystem models under RCP2.6 and RCP8.5 without fishing for different size groups (n=10). Box plots display the median (horizontal line), mean (x) and interquartile range (boxes).

The magnitude and variability of the climate-change effect (RCP8.5 vs. RCP2.6) were comparable between a fished and unfished ocean (Fig. 1d, Fig. S3a), suggesting that fishing, at least under similar fishing pressure to present, may not substantially alter the relative effect of climate change. However, the mean climate effect was slightly weaker in a fished ocean (Fig. 1d, Fig. S3a), possibly due to lower biomass levels and temperature-dependent effects^9^. Warming enhances both growth and predation rates, yet in a fished ocean predation rates are reduced due to selective fishing and lower predator abundance^16-17^, thereby mitigating biomass changes^9^. This is a relatively small effect, and pales beside the magnitude of biomass decrease due to fishing itself (Table S2). We note, however, that the magnitudes of the fishing effects are not directly comparable due to differences in how fishing pressure and commercial taxa are incorporated in the different EMs. We also caution that our constant future fishing scenario is simplistic and does not incorporate changes in effort, technology, management and conservation^9,18-20^, which could substantially increase or decrease fishing pressure and modify projected biomass trends.

Although ensemble means revealed consistent global biomass declines across different scenarios, there was considerable variation among individual EM-ESM-combinations (Fig. 1b, Fig. S2). Generally, projected changes based on GFDL-ESM2M were lower than those based on IPSL-CM5A-LR, reflecting lower SST increases and smaller NPP reductions in GFDL-ESM2M (Fig. S1). This reinforces previous work highlighting the importance of ESM uncertainty in future projections of fish biomass and potential fisheries production^21-22^. Interestingly, the variability in projected biomass changes among different ESMs was of similar magnitude as that among different EMs (Fig. 1e), suggesting similar levels of uncertainty in both modeling fields. Other MIPs also found that uncertainties in both ESMs and climate-impact models contribute to overall projection uncertainty, with similar contributions from ESMs and global hydrological models^13^, yet much higher uncertainty in global crop and vegetation models^14-15^.

Variability among EMs can be attributed to differences in fundamental structures, species or functional groups included, and ecological processes incorporated (Table 1, Table S1). Thus, some models strongly respond to changes in temperature affecting metabolic rates (BOATS, Macroecological), while others are largely driven by changes in NPP affecting trophic dynamics (EcoOcean), or a combination of temperature, NPP, and additional drivers (e.g. oxygen, pH, ice cover) affecting habitat niches and species distribution (DBEM) or overall food-web dynamics (DPBM, APECOSM)^6-11^. Notably, the variability of projected changes among both EMs and ESMs was higher under RCP8.5 than RCP2.6 (Fig. 1e), and that among EMs higher in animals >30cm (Fig. 1e, Tables S2), suggesting greater uncertainty of projections with stronger warming and in larger animals.

Many policy-makers and processes use the change in global air temperature since pre-industrial times as a reference for the effects of climate change^23-24^. For our ensemble projections, this revealed a predictable linear impact with an average 5%-drop in total animal biomass with every 1ºC of Earth-surface warming in the absence of fishing (Fig. 2). Similar declines were found for animals >10cm and >30cm (Fig. S4). These relationships may be slightly less negative in a fished ocean (Fig. S3b), given the dampening effect described above, yet remain substantial. These model-derived relationships are simplistic, and natural ecosystems show likely more complex responses to warming^1,25^, but may be useful approximations for the global average. According to our results, limiting global warming to 1.5-2ºC above pre-industrial levels would lead to a 4-6% decline in the capacity of the global ocean to sustain marine animal biomass (Fig. 2), underscoring the potential impacts of mitigation measures in accordance with the Paris Agreement^23-24^.

**Figure 2.**
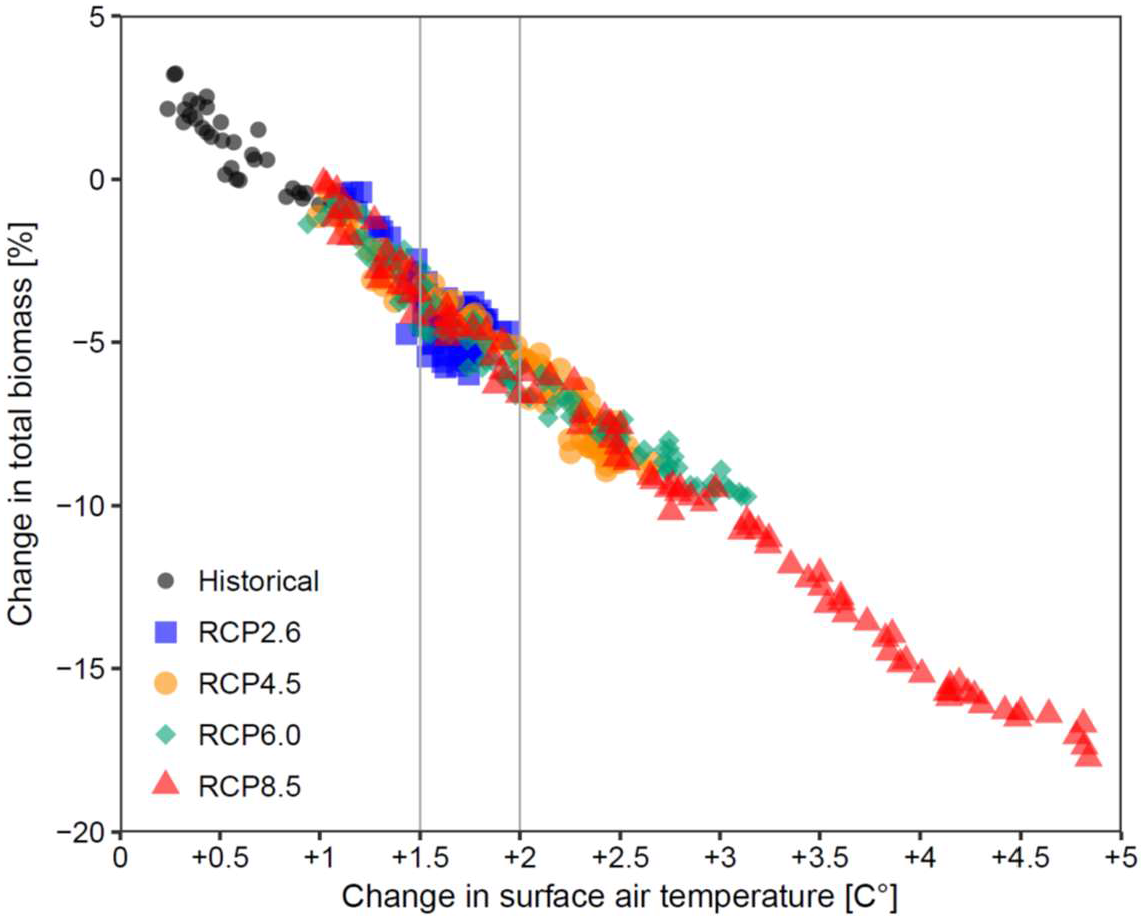
Relationship of the change in marine animal biomass with increasing global surface air temperature since pre-industrial times (1860s). Each dot represents an annual ensemble mean (n=10) relative to 1990-99 across historical and future emission scenarios (RCPs) in the absence of fishing. Vertical lines frame expected changes between 1.5° and 2°C warming.

Not all regions in the global ocean respond similarly to changing climates (Fig. 3, Fig. S5). Our ensemble projections revealed strong increases in total animal biomass in polar regions and widespread declines in temperate to tropical regions under RCP8.5 (Fig. 3b), with qualitatively similar but much less pronounced patterns under RCP2.6 (Fig. 3a), again highlighting the benefits of climate-change mitigation. These climate-change effects were spatially similar in a fished and unfished ocean (Fig. S6). However, our ensemble projections differed spatially from previous single-model results highlighted in the IPCC’s Fifth Assessment Report^3^; particularly, we found strong biomass declines (not increases^3^) in temperate to subtropical regions and increases (not declines^3^) around Antarctica. Our spatial patterns were more in line with those projected for marine pelagic communities^10^, whereas the magnitude of global and regional changes varied from other single-model results^6-9^. This points to the value of ensemble projections and model inter-comparisons and highlights the risk of relying on single-model results. While warming waters and enhanced primary production facilitate species expansions and biomass increase in polar regions, tropical areas may experience pronounced species losses as thermal thresholds are exceeded^1^. In temperate regions, warming is expected to change species composition and, coupled with enhanced water-column stratification reducing primary production, declines in animal biomass^1^.

**Figure 3.**
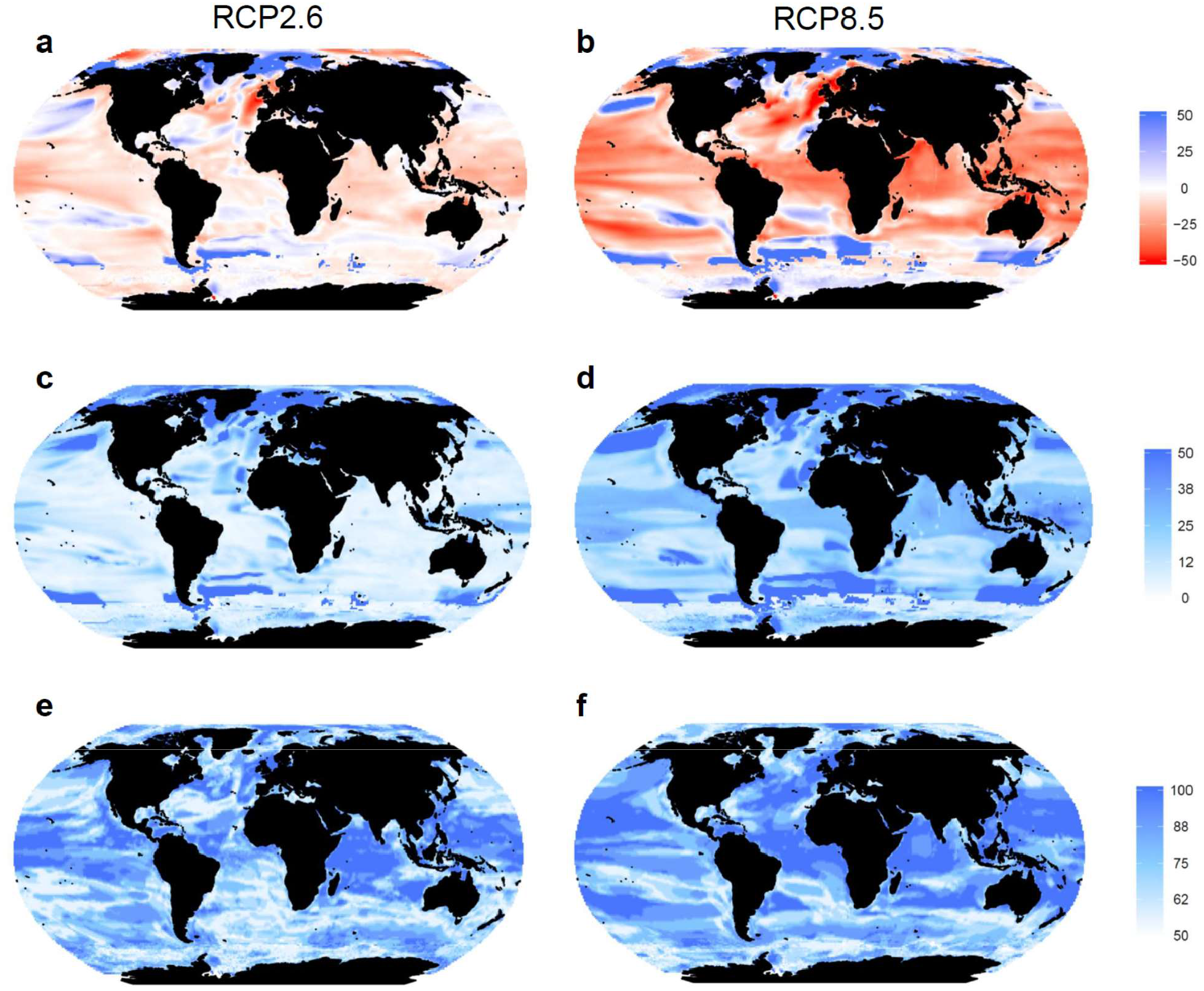
Global spatial patterns of ensemble projection results for RCP2.6 (left) and RCP8.5 (right). (**a, b**) Multi-model mean change (%, n=10) in total marine animal biomass in 2090-99 relative to 1990-99 without fishing. (**c, d**) Variability among different ecosystem and Earth-system model combinations expressed as one standard deviation. (**e, f**) Model agreement (%) on the direction of change.

The combination of different ecosystem structures and processes represented in our ensemble projections more likely reflects future trends and their uncertainties than any single-model result, particularly in regions with high model agreement on the direction of change (75-100%; Fig. 3e-f). The magnitude of change, however, varied with relatively low inter-model variability in temperate to tropical regions, but considerable variability in many polar and coastal regions (Fig. 3c-d). Again, some of this variability is due to differences among ESMs (Fig. S7) and some due to differences among EMs (Fig. S8). One major caveat is that while ESMs capture climate variations across major ocean biomes, their coarse resolution limits representation of some coastal and polar processes^26^. Similarly, global EMs do not resolve diverse coastal habitats, such as coral reefs or kelp forests, nor their contributions to primary production and biogeochemical cycles. Thus, projections based on global models are less reliable in coastal and polar regions but could be improved through finer-scale global models or coordinated regional downscaling to incorporate higher resolution climate and ecosystem features^5,26^. Also, some regional models may more accurately capture processes at the scales at which most management occurs^27^. Here, a MIP comparing regional models to their respective regions extracted from global models may prove as a useful next step^11^, as well as finer-scale ensemble projections on climate-change effects on fish stock^18,28^.

By providing estimates of global and regional biomass changes and associated uncertainties, our ensemble projections represent the most comprehensive outlook on the future of global marine animal biomass to date, and highlight the impacts and risks associated with differing climate-change scenarios. Our results are robust in terms of the direction of change, yet the significant spread in the magnitude of projections illustrates considerable variation due to uncertainty in both ESMs and EMs. The challenge will now be to understand and address these uncertainties to improve our ability to predict the response of marine ecosystems to climate change on different temporal, spatial and ecological scales. The next round of climate-model projections (CMIP6) with improved and finer-scale representation of biogeochemical parameters that drive many key processes in marine ecosystems will likely advance future ensemble projections^11^. The incorporation of additional EMs based on novel paradigms or reflecting alternative structures and processes may also be informative^11^, and the comparison between regional- and global-scale models^11,27^. Future EMs characterizing individual species also need to better resolve the capacity of marine organisms for acclimation and adaptation to rapidly changing environments, and the extent to which these responses modify projected distribution and abundance. In size-based EMs, species composition is not resolved, and projections are not expected to be sensitive to the effects of phenotypic and evolutionary change, because the broad relationships between temperature, metabolic rate and body size still apply as food webs are reconfigured. Further, the continued expansion of global observational datasets provides opportunities to better constrain and validate global models. Finally, a large component of future change will depend on the trajectories of fisheries, aquaculture and other human impacts in the ocean, and the demand for food and feed on land^4,9,18,29^. Over the past century, fishing has caused much stronger biomass declines^16-17^ than the projected climate-change effects presented here. Incorporating standardized temporally- and spatially-resolved scenarios of human activities and alternative management approaches will be crucial to improve our understanding of the future of marine fish and ocean ecosystems^11,20^ and identify points of greatest leverage for mitigating impacts.

Our results highlight that, by 2100, global marine animal biomass is expected to decline by 17% under business-as-usual greenhouse gas emissions (RCP8.5) without fishing, suggesting substantial climate-driven changes to ocean ecosystems and challenges for human society. Meeting the SDGs on food security (SDG2), livelihoods (SDG1) and well-being (SDG3) for a growing human population by 2030 while also sustaining life below water (SDG14) represents a near-term example of this challenge. Our ensemble projections indicate greatest decreases in future animal biomass at mid-low latitudes, where many nations strongly depend on seafood and fisheries, and where marine biodiversity is already threatened by multiple human activities^4,29^. In turn, greatest increases are projected at high latitudes, highlighting new opportunities for (and conflict over) resource use, but also chances for protecting sensitive species and rapidly changing polar ecosystems. Improved dynamic and adaptive fisheries and ecosystem management and conservation may mitigate some future climate-change impacts and maintain ecosystem health and service provision^4,17-19^. However, this can only happen if the international community, including national and regional bodies strengthen the required institutions and management approaches^4,17,30^.

## Methods

### Ecosystem model selection

We used six published and peer-reviewed global fisheries and marine ecosystem models (EMs) that participated in the first round of the Fisheries and Marine Ecosystem Model Inter-comparison Project (Fish-MIP)^11^. The six EMs varied considerably in basic model structure and underlying assumptions, taxonomic scope and key ecological processes included, and their representation of species, functional groups, size classes and the proportion of commercial to non-commercial taxa (Table 1, Table S1). Together, this heterogeneity reflects the diversity of model structures, parameterizations, scopes and purposes, meaning that our ensemble is more likely to include a greater number of relevant processes in the ocean than any single model.

The EMs also varied in their representation of fishing, with some models only representing an unfished ocean (Macroecological) while others able to incorporate fishing effort (BOATS, EcoOcean) or fishing mortality (DBEM, DPBM, APECOSM) as forcing variables, but so far only few models can incorporate feedbacks from the biological to social and economic systems. For example, BOATS uses an interactive bioeconomic model to determine spatial and temporal changes in fishing effort, and EcoOcean uses a spatially-explicit fishing dynamics model to simulate spatial patterns of imposed fishing effort^9,11^. Due to these inherent differences, we could not standardize fishing scenarios across EMs and therefore assessed the climate-change effect in an unfished compared to a fished ocean in a subset of EMs (see below).

### Climate model selection

All EMs were forced with the same standardized set of outputs from ESMs derived from the Coupled Model Inter-comparison Project Phase 5 (CMIP5 database: http://cmip-pcmdi.llnl.gov/cmip5/). We selected NOAA’s Geophysical Fluid Dynamics Laboratory Climate Model (GFDL-ESM2M)^31^ and the Institute Pierre Simon Laplace Climate Model (IPSL-CM5A-LR)^32^ because they generated all necessary physical and biogeochemical outputs needed to run our global marine EMs, particularly the monthly depth-resolved biogeochemical fields of different size groups of phytoplankton and zooplankton concentration and productivity^11^. While several other ESMs are available in CMIP5, they did not generate or save all variables required by our EMs. The two selected ESMs spanned the range of projections from all CMIP5 models with GFDL-ESM2M being at the lower end and IPSL-CM5A-LR at the higher end of projected future changes in SST and NPP (Fig. S1), while other variables (e.g. pH, O_2_ concentrations) were more similar among all CMIP5 models^12^. Therefore, global annual mean trends of these two ESMs should reflect the multi-model mean of all CMIP5 models^12^.

Each ESM was run for each of four Representative Concentration Pathways (RCPs) representing a standard set of IPCC informed emission scenarios, ensuring that starting conditions, historical trends, and projections are consistent across models and sectors. RCP2.6 represents a strong mitigation scenario characterized by an emission pathway leading to very low greenhouse gas (GHG) levels by 2100^33^. RCP4.5 and RCP6.0 represent stabilization emission scenarios, requiring a stabilization in radiative forcing after 2100, without exceeding the target value of 4.5 W m^-2^ for RCP4.5 and 6.0 W m^-2^ for RCP6.0^34^. RCP8.5 represents a business**-**as**-**usual scenario characterized by increasing GHG emissions over time leading to high GHG emissions in 2100^35^.

### Standardized model inputs and outputs

The fundamental goal of Fish-MIP is to compare the response of a wide range of marine EMs to common external forcings, namely standardized climate-change scenarios. Standardized physical and biogeochemical variables from the 2 ESMs and 4 RCPs were derived on a 1×1 global grid for a historical (1970-2005) and a future period (2006-2100) to be used as climate input data in our EMs. Physical variables included current velocities, water temperature, salinity, dissolved oxygen concentration, pH, mixed-layer depth and ice coverage. Biogeochemical variables included large and small phytoplankton and zooplankton concentrations and productivity. All variables were summarized as monthly, depth-resolved values and then further aggregated or integrated to suit the input requirements for each EM, such as providing values for surface, bottom or depth-integrated layers. For more details on input variable selection and requirements of each ecosystem model see ref. 11.

First, we used a no-fishing scenario across all six EMs to isolate the climate-change effect on fish or animal biomass. In a subset of three EMs we also used a simple fishing scenario based on observed or estimated time-varying data on fishing effort, mortality or exploitation rates (depending on model requirements) for the historical period (1970-2005) and then kept constant at 2005 levels for the future period (2005-2100)^11^. Thereby, each model relied on its own mechanism for incorporating fishing data^11^. Dynamic projections of future fishing pressure were not used because appropriate temporally- and spatially-explicit scenarios of future fishing pressure are not yet available^11^, although we do acknowledge that fishing pressure will change over the projection period. Recently, future fishing scenarios have been developed conceptually based on the IPCC’s Shared Socio-economic Pathways (SSPs)^20^. Once translated into quantitative form, these should be available for future efforts to project changes in fish and fisheries.

All EMs were required to produce a set of standardized outputs that could be directly compared and combined into ensemble projections^11^. For this study, we selected three outputs:(1) total consumer biomass density (g C m^-2^) representing all animals, size classes or trophic groups in each model, (2) biomass density of all animals, size classes or trophic groups >10cm, and (3) biomass density of all animals, size classes or trophic groups >30cm. Since the taxonomic scope differs among models, we used the terms ‘total animal biomass’, ‘animals >10cm’ and ‘animals >30cm’, respectively, for simplification.

### Simulations

Several limitations resulted in not all EMs being able to run the full set of 16 simulations from 2 ESMs, 4 RCPs and 2 fishing scenarios for total animal biomass, biomass of animals >10 cm and >30 cm. We therefore decided to use comparable subsets of available EMESM combinations to answer different questions. ESM limitations included the unavailable monthly, depth- and size-resolved biogeochemical data in GFDL-ESM2M (not run in APECOSM, DPBM); EM limitations included substantial simulation run time in some EMs (e.g. DBEM did not run all RCPs; DPBM and APECOSM did not perform fishing runs), the lack of size class differentiation (e.g. DBEM), and the inability to incorporate fishing (e.g. Macroecological). Therefore, all six EMs used the IPSL-CM5A-LR no-fishing runs and four used the GFDL-ESM2M no-fishing runs for RCP2.6 and 8.5 (n=10), and five and three EMs, respectively, for RCP4.5 and 6.0 (n=8; Table S2). A subset of three EMs (BOATS, EcoOcean, DBEM) was used for fishing runs with inputs from both ESMs for RCP2.6 and 8.5 (n=6). All six EMs projected total animal biomass, and five EMs animal biomass >10cm and >30cm (Table S2). However, since DBEM represents all commercial fish and invertebrate species covering the full size spectrum, we used DBEM outputs for all three size groups, but cross-checked results with and without DBEM as a sensitivity analysis.

### Analyses

Because the six EMs include different species, size classes and trophic groups, their output for absolute biomass density varied across models. We therefore calculated time series of relative change in animal biomass defined as percent (%) relative to 1990-99 for each simulation. This reference period was chosen as representing the last decade of the 20th century, which was later compared to 2090-99 as the last decade of the 21st century. The different time series of relative change were then averaged into a multi-model mean change, and variability among models was described with the standard deviation around the mean (Fig. 1a, Fig. S2) as well as with box and whisker plots (Fig. 1c). To summarize the results across simulations and size groups, we also calculated the % change in 2090-99 relative to 1990-99 (Table S2).

Given the differences in EM structure and characterization of fishing (see above) we could not standardize the fishing scenarios and did not compare the magnitude of the fishing effects across models. Instead we compared the relative difference in the climate effect (RCP8.5 vs RCP 2.6) in a fished and an unfished ocean. To do so, we calculated ((RCP8.5-RCP2.6)/RCP2.6) within each EM in the 2090s and over time which was then compared across EMs (Fig. 1d, Fig. S3a).

Next, we compared the variability of results due to the different ESMs and EMs (Fig. 1e). For ESM variability, we calculated the standard deviation for individual EM results for simulations run with the IPSL-CM5A-LR compared to the GFDL-ESM2M (for all EMs run with both forcings, n=4). For EM variability, we calculated the standard deviation of results across all EMs for i) only the IPSL-CM5A-LR and ii) only the GFDL-ESM2M. These calculations were performed separately for the different RCPs and size groups in an unfished ocean (Fig. 1e).

To relate the magnitude of change in total animal biomass (Fig. 2) and biomass of animals >10cm and >30 (Fig. S4) to changes in global air temperature since pre-industrial times, we derived global air temperature data from 1861-2100 for both ESMs. We calculated the average pre-industrial temperature for the period 1861-1870, and then the relative change in air temperature over this reference period. We then plotted the % change in total animal biomass over the change in air temperature across the historical and future period including all RCPs in an unfished ocean (Fig. 2). This was repeated for animals >10cm and >30cm (Fig. S4). To assess how this relationship might change in a fished ocean, we plotted the difference in % biomass change (RCP8.5 vs RCP2.6) in a fished versus unfished ocean over the difference in global surface air temperature between RCP8.5 vs RCP2.6 in any given year (Fig. S3b). The resulting positive relationship suggest that as warming increases, the reduction in biomass is lesser in a fished compared to an unfished ocean.

To visualize spatial patterns of change globally, we mapped the relative (%) change in 2090-99 compared to 1990-99 for the multi-model mean and standard deviation of total animal biomass on a 1×1 degree grid across all EM-ESM-combinations for RCP 2.6 and RCP8.5 in an unfished ocean (Fig. 3). We also mapped the multi-model mean +1SD and −1SD (Fig. S5) to visually display the full range of minimum to maximum potential changes. As an additional measure of robustness, we mapped the percentage of EMs agreeing on the direction of change^12^, where 100% means full model agreement and 50% represents an even split. Next, to compare the climate effect in a fished and an unfished ocean spatially (Fig. S6), we mapped the multi-model mean relative difference (%) in biomass change (RCP8.5 vs RCP2.6) in 2090-2099 in a fished and in an unfished ocean based on the three EMs that performed fishing runs (BOATS, DBEM, EcoOcean). To evaluate the spatial variability due to ESM selection in an unfished ocean (Fig. S7), we also mapped the multi-model mean biomass change and standard deviation across all EMs for the IPSL-CM5A-LR and the GFDL-ESM2M separately. Lastly, we mapped relative changes in total animal biomass for each EM for IPSL-CM5A-LR and GFDL-ESM2M under RCP2.6 and RCP8.5 in an unfished ocean (Fig. S8).

### Supplementary information

Table S1-S2, Figures S1-S8

## Acknowledgments

We thank B. Worm for valuable discussions and comments. Financial support was provided by the German Federal Ministry of Education and Research (BMBF, grant no. 01LS1201A1) through the Inter-Sectoral Impact Model Intercomparison Project (ISI-MIP), and the European Union’s Horizon CERES project (grant no. 678193). HKL and WWLC acknowledge financial support from the Natural Sciences and Engineering Research Council (NSERC) of Canada, DPT from the Kanne Rassmussen Foundation Denmark, ABB from the NSERC CREATE Transatlantic Ocean Science and Technology Program, WWLC and TE from the Nippon Foundation-UBC Nereus Program, EDG from the European Research Council (grant no. 682602), MC and JS from the European Union’s Horizon MERCES project (grant no. 689518), EAF, JLB and TR from CSIRO and the Australian Research Council (Discovery projects DP140101377, DP170104240), NB, LB and OM from the Agence Nationale de la Recherche (grant no. ANR-17-CE32-0008-01) and the Pôle de Calcul et de Données Marines (PCDM) for providing DATARMOR storage, data access and computational resources, and SJ from the UK Department of Environment, Food and Rural Affairs.

